# RVAgene: Generative modeling of gene expression time series data

**DOI:** 10.1101/2020.11.10.375436

**Authors:** Raktim Mitra, Adam L. MacLean

## Abstract

Methods to model dynamic changes in gene expression at a genome-wide level are not currently sufficient for large (temporally rich or single-cell) datasets. Variational autoencoders offer means to characterize large datasets and have been used effectively to characterize features of single-cell datasets. Here we extend these methods for use with gene expression time series data. We present RVAgene: a recurrent variational autoencoder to model gene expression dynamics. RVAgene learns to accurately and efficiently reconstruct temporal gene profiles. It also learns a low dimensional representation of the data via a recurrent encoder network that can be used for biological feature discovery, and can generate new gene expression data by sampling from the latent space. We test RVAgene on simulated and real biological datasets, including embryonic stem cell differentiation and kidney injury response dynamics. In all cases, RVAgene accurately reconstructed complex gene expression temporal profiles. Via cross validation, we show that a low-error latent space representation can be learnt using only a fraction of the data. Through clustering and gene ontology term enrichment analysis on the latent space, we demonstrate the potential of RVAgene for unsupervised discovery. In particular, RVAgene identifies new programs of shared gene regulation of *Lox* family genes in response to kidney injury.

## 1 Introduction

Dynamic changes in gene expression control the transcriptional state of a cell, and are responsible for modulating cellular states and fates. Gene expression dynamics are in turn controlled by cell-internal and external signaling networks. Despite the noisiness of gene expression in single cells (Raj & Van Oudenaarden 2008), over time or over populations of cells, predictable patterns emerge. Here we address the challenge of classifying and predicting gene expression dynamics across large groups of genes.

Machine learning (and deep learning in particular) has led to recent advances in our ability to explain or predict biological phenomena (Ching et al. 2018). Deep learning modeling via autoencoders (Hinton & Salakhutdinov 2006) and variational autoencoders (Kingma & Welling 2014) has been central to progress in the field. Autoencoders learn two functions: one to encode each input data point to a low dimensional point, and another (the decoder) to reconstruct the original data point from the low dimensional representation. Variational autoencoders (VAEs) build on this architecture and instead encode input data points as distributions; VAEs are less prone to overfitting and can offer meaningful representations of biological features in the latent space (Way & Greene 2017).

Single-cell mRNA sequencing (scRNA-seq) data present appealing sources of data for deep learning models, given their size and complexity (Svensson et al. 2018). Deep learning models have been used to analyze scRNA-seq data and address a variety of challenges. Autoencoders have been developed to perform noise removal/batch correction (Deng et al. 2019, Eraslan et al. 2019, Wang et al. 2019), imputation (Talwar et al. 2018), and visualization & clustering (Lin et al. 2017). VAEs have been developed for the visualization and clustering of scRNA-seq data (Ding et al. 2018, Wang & Gu 2018), and can provide a broad framework for generative modeling of scRNA-seq data (Lopez et al. 2018): scVI can be used for batch correction, clustering, visualization, and differential expression testing.

The methods described above for single-cell data analysis by deep learning focus primarily on cell-centric tasks; here we are interested in gene-centric inference. Particularly, we are interested in characterizing dynamic changes in gene expression. These can be either changes with respect to real time or “pseudotime,” the latter referring to the ordering of single cells along an axis describing a dynamic cell process such as development or stem cell differentiation (see methods overview in (Saelens et al. 2019)). We can interpret any scRNA-seq data as gene expression time series data, given an appropriate underlying temporal process, either in terms of real (experimental) time (low resolution: around 2 - 20 data points) or pseudotime (high resolution: 10^3^ - 10^6^ data points). McDowell et al. (2018) introduced a non-parametric hierarchical Bayesian method (DPGP) to model such data. Using a Gaussian process to cluster temporal gene profiles and a Dirichlet process to generate the Gaussian processes, DPGP offers powerful and intuitive means with which to cluster gene expression time series data. However, since learning Gaussian processes is equivalent to a fully agnostic search in function space, training DPGP is computationally intensive and difficult to parallelize.

Clustering relies on strong assumptions about the underlying structure of the data. Even for methods that move away from hard clustering towards probabilistic methods for cell type assignment (Jetka et al. 2018, Zhu et al. 2019), assumptions remain and under certain conditions a continuous representation of the data may be better. Here we take such an approach, and seek to find a low dimensional representation of the data, on which further analyses (including but not limited to clustering) can be performed. VAEs are an obvious choice, given their success on other scRNA-seq analysis tasks, but modeling temporal changes with a feed-forward VAE would be equivalent to a fully agnostic search, similar to learning a Gaussian process. Recurrent networks offer well-established architectures for learning sequential and temporal data, and have been successfully combined with VAEs (Fabius & van Amersfoort 2014). We use a recurrent network architecture to structure the data and reduce the computational cost.

We introduce a recurrent variational autoencoder for modeling gene dynamics from scRNA-seq data (RVAgene). RVAgene learns two functions during training, parameterized by encoder and decoder networks. The encoder network projects the training data into latent space (we use a 2 or 3 dimensions in order to visualize, though there are no inherent limits). The decoder network learns a reconstruction of training genes from their latent representation. RVAgene facilitates clustering of other characterization of gene profiles in the latent space. By sampling points from the latent space and decoding them, RVAgene provides means to generate new gene expression time series data, drawn from the biological process that was encoded. Overall, RVAgene serves as a multipurpose generative model for exploring gene expression time-series data.

The remainder of the paper is structured as follows: we next present methodological details and development of RVAgene. We produce a synthetic gene expression time-series dataset with innate cluster structure, and demonstrate the accuracy of RVAgene on these data. We then explore two biological datasets with RVAgene: a scRNA-seq dataset on stem cell differentiation, where gene expression changes are studied over pseudotime; and a bulk RNA-seq dataset describing dynamic responses to kidney injury. We conclude by discussing key features and limitations of RVAgene, and recent advances in machine learning that will pave the way for future work in these directions.

## 2 Methods

We develop a recurrent variational autoencoder to model gene expression dynamics (RVAgene). Here we briefly describe the methods underpinning variational autoencoders, and present the implementation of RVAgene.

### 2.1 Variational inference and variational autoencoders

In the most general setting of a Bayesian model, we seek to learn the latent variables **z** that best characterize some data **x**. Given a generative process that draws latent variables from a prior distribution, *p*(**z**), and a likelihood of the data observed that is given by *p*(**x**|**z**), then the posterior probability is given by Bayes rule:

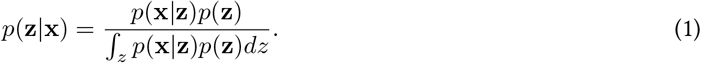

The denominator is often intractable, making it difficult to estimate *p*(**z|x**). Markov Chain Monte Carlo methods provide means to estimate posterior probability distributions. An alternative method to estimate hard-to-compute probability distributions is Variational Inference (VI) (Hoffman et al. 2013), which starts from the assumption that the posterior can be approximated by a distribution *q*(**z**) from the family 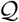. VI then amounts to an optimization problem to find the *q*\* that minimizes the Kullback–Leibler (KL) divergence between the approximation and the true posterior:

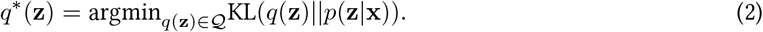

Much recent effort has gone into solving VI problems in different settings (Zhang et al. 2018, Ingraham & Marks 2017, Bouchard-Côté & Jordan 2010). VI can be framed as solving an optimization problem over function families: neural networks are popular candidates for representing and learning complex functions. VI was incorporated into autoencoders (Kingma & Welling 2014) to create the architecture of a variational autoencoder (VAE). A VAE consists of an encoder network to approximate *p*(**z|x**) through a function *q***x**(**z**), and a decoder network *p*(**x|z**) (Fig. 1A). Conceptually, the encoder solves an inference problem: approximating the posterior distribution *p*(**z|x**) as some 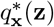, while the decoder solves a reconstruction problem: defining a generative process for *p*(**x|z**), given the latent variables. The VAE posterior is modeled by a multivariate normal 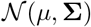 of the same dimension as **z**. Training then comes down to minimizing two objective functions. For the encoder network, which should learn a “well distributed” latent space, minimize the KL divergence: 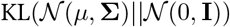. For the decoder network, which should reconstruct the inputs **x** from the latent space, minimizing either an *L*1 or *L*2 objective function with respect to 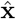 is appropriate. The use of KL-divergence and an *L*2 objective solves the VI formulation of Eq. 2 (Kingma & Welling 2014), however, an *L*1 objective may be preferred in practice, e.g. in cases where we want to suppress the effects of outliers on the structure of **z**(Botchkarev 2018).

**Figure 1.**
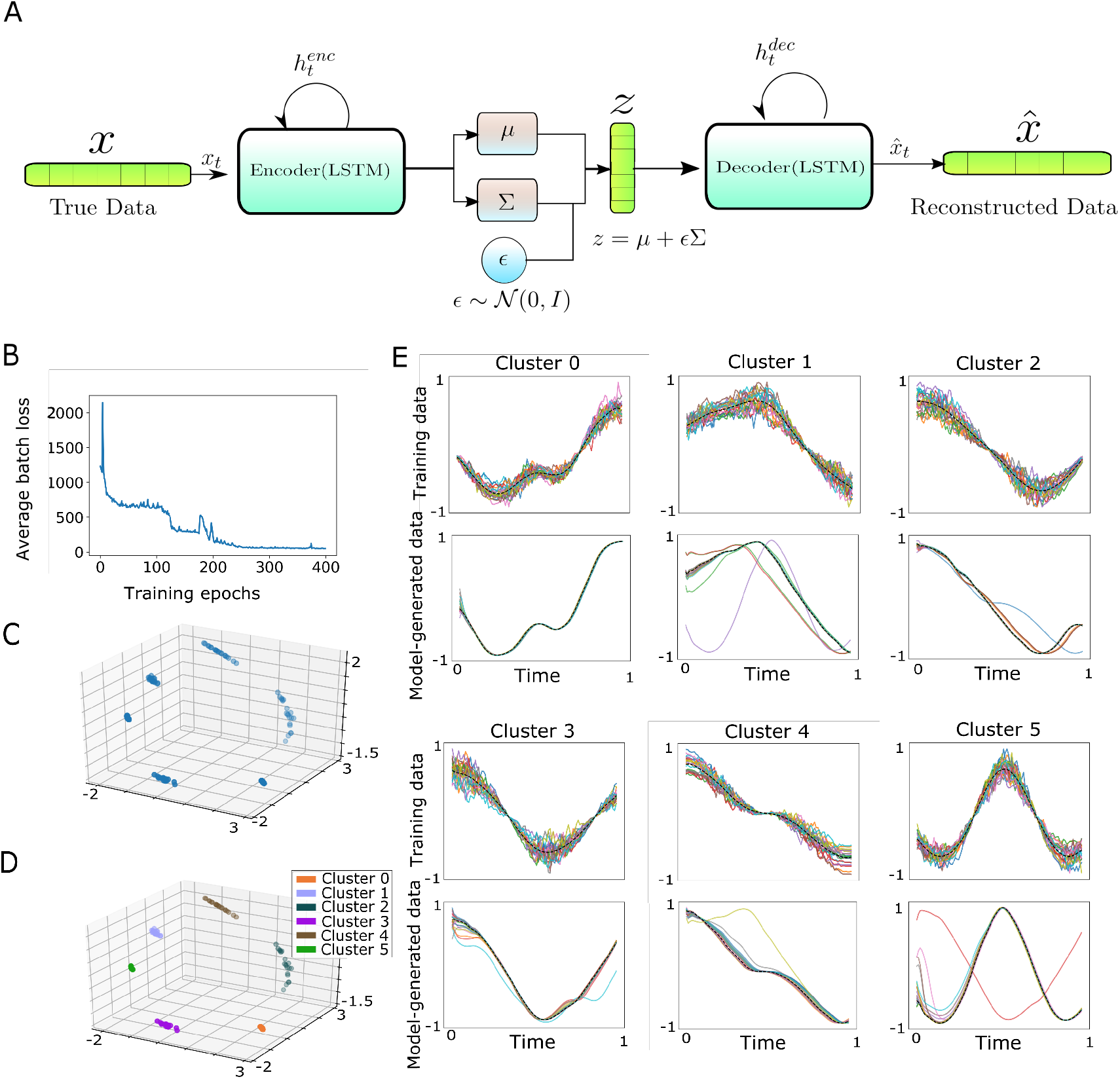
Unsupervised representation learning with RVAgene using synthetic data. **(A)** Schematic diagram of the RVAgene model. (**B**) Average loss function 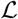 as over duration of training. (**C**) Latent space representation learnt by RVAgene model after training. (**D**) Clusters detected by *k*-means clustering on the latent space, with *k* = 6 (**E**) First and third rows show input training data used (20 simulated genes in each of six clusters); cluster means shown in black. Second and fourth rows show the model-generated data, obtained by sampling and decoding points from the latent space; decoded cluster empirical means shown in black.

### 2.2 RVAgene: A recurrent variational autoencoder to model gene expression dynamics

Following the VAE architecture, RVAgene consists of an encoder and a decoder network with a reparameterization step in between. To incorporate the knowledge that we are modeling temporal data, recurrent neural networks offer an ideal architecture to use for both the encoder and the decoder networks. Recurrent and VAE networks have been successfully combined elsewhere, e.g. for textual (Nallapati et al. 2016) and time series data (Malhotra et al. 2015).

The architecture of RVAgene is based on Fabius & van Amersfoort (2014). An input sequence (i.e. gene) *x* ∈ **x**, *x* = (*x*_1_, *x*_2_,…, *x_t_*,…, *x_T_*) is encoded using a recurrent function described by a long short-term memory (LSTM) unit. LSTM units are the state-of-the-art in recurrent architectures, since they are robust against the vanishing gradient problem for longer sequences, unlike other recurrent units (see details in Hochreiter & Schmidhuber (1997)). We encode *x* in the following manner:

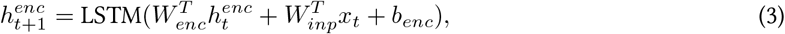

where (*W_enc_*, *W_inp_* and *b_enc_*) are network weight parameters, and the hidden states *h_t_* represent information shared over timepoints in the LSTM. The dimension of the *h_t_* (and *W_enc_*) is given by a hyperparameter (“hidden-size”). The encoded *h*_*t*+1_ are used to parametrize the posterior mean and variance from *x*, with mean *μ_z_* and diagonal covariance *σ_z_* as:

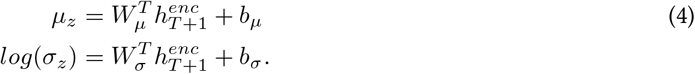

We then use the reparameterization trick described in Kingma & Welling (2014) to sample *z* from the distribution:

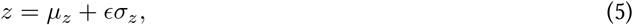

where, for known *ϵ*, backpropagation through the sampling step is possible while training the network.

For the decoder network, the first state *h*_1_ is calculated from *z*, and the recurrent formulation follows by reconstructing *x* as 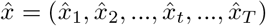, thus:

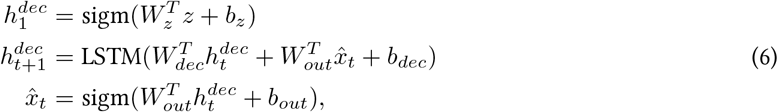

where 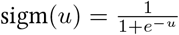 is the sigmoid activation function, and (*W_i_, b_i_*) are the network weight parameters. A schematic diagram of the network is shown in Fig. 1A, which can now be trained using backpropagation, to minimize the objective function:

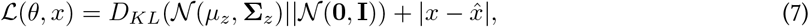

where *μ_z_* and **Σ**_*z*_ = diag(*σ_z_*) are calculated from *x* by the encoder.

To evaluate the accuracy of RVAgene, we need an appropriate error measure. For each gene in the test set, we calculate the *L*1 reconstruction error between generated data 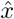 and true data *x*, averaged over all time points. We normalize the data to lie in [0, 1] to avoid skewing the error by differences in gene expression magnitudes. Thus we define:

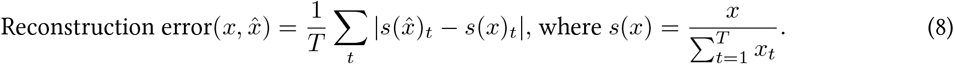

### 2.3 Generating synthetic gene expression time series data

To test RVAgene, we generate a synthetic time series dataset. Six clusters each containing 20 genes are simulated, where for each cluster *c*, the mean gene expression time series *Y_c_* = (*y*_*c*1_, *y*_*c*2_,…, *y_ct_*) was generated using addition or convolution and rescaling of two random sinusoidal functions of the form *k*_1_sin(*k*_2_*t*), where *k*_1_, *k*_2_ are randomly chosen positive integers. Trajectories of cluster members were then generated by sampling from the multivariate normal 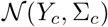. We model Σ_*c*_ as the positive definite matrix 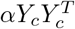, where *α* is a scaling factor, we use: 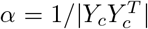. As defined, Σ_*c*_ will describe nonzero correlations for all pairs of time points, (*t_i_, t_j_*). This is unrealistic, so we set to 0 the entries of Σ_*c*_ for which column and row indices have a difference of more than some threshold *T* (we used *T* = 50), reflecting the fact that correlations between time points are lost over larger time windows (temporal correlations are local). Note that under this condition, Σ_*c*_ is no longer necessarily positive definite. The multivariate Gaussian sampler `numpy.random.multivariate_normal()` implemented in numpy (Harris et al. 2020) was used to sample from this augmented Σ_*c*_.

## 3 Results

### 3.1 RVAgene can accurately and efficiently reconstruct temporal profiles from synthetic data

We generated a dataset of 120 genes using convolutions of sinusoidal functions (see Methods) to test the ability of RVAgene (Fig. 1A) to learn and predict noisy nonlinear temporal profiles. An RVAgene model was trained on all 120 genes from 6 clusters with a hidden size of 70 and a 3 dimensional latent space. The model was trained for 400 epochs, after which the average batch objective 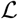 function indicates convergence (Fig. 1B), producing a three-dimensional latent space representation (Fig. 1C). K-means clustering on the latent space (k=6) identified well-separated clusters (Fig. 1D).

RVAgene modeling followed by k-means clustering on the latent space identified 6 clusters with perfect fidelity between predicted and true clusters. One might reasonably ask, why use a neural network for this task? Simpler dimensionality reduction methods (e.g. PCA, t-SNE, or a non-variational autoencoder) would also find the correct solution. RVAgene has the advantage over these methods that the underlying structure of the latent space leads to interpretability. A point in reduced PCA or t-SNE space that does not overlap with a data point is not interpretable. Traditional autoencoders lack regularity in the latent space, i.e. even for a representation with arbitrary accuracy (a reconstruction error of zero), decoding a point that does not correspond to a training data point can result in nonsensical generated data, even if the decoded point is arbitrarily close to a training data point. Variational Auoencoders remedy this by learning a regularised or smoother distribution on the latent space. In this sense, the KL-divergence term in the VAE loss function can be thought of as a regulariser. This property enables RVAgene to generate new gene expression dynamics by decoding points from different regions of the latent space, having properties similar to clusters nearby to those points.

To demonstrate the generative properties of the RVAgene latent space, we sample points from multivariate Normal distributions, centered on the empirical mean of each cluster with variance of 0.4, i.e. 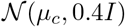, where *μ_c_* is the empirical mean of the cluster and **I** is the identity matrix in ℝ^3^. Corresponding to each cluster, we sample 20 points in the latent space, and use the decoder network to generate new time series data (Fig. 1E). Most of the points sampled generate trajectories that belong to the correct cluster. Moreover, we identify cases where corresponding to transitions between clusters. For example, some points sampled near Cluster 2 generate trajectories that are similar to members of Cluster 4, and vice versa. This makes sense due to the similarity between the temporal profiles of Clusters 2 and 4. A similar correspondence is observed between Clusters 1 and 5. We also observe some generated trajectories that display intermediate profiles between two or more clusters: the decoder function learnt by RVAgene is smooth, and gives rise to meaningful representations of points across regions of the latent space.

RVAgene offers additional functionality as a tool for removing noise the data. RVAgene model reconstructions generated by decoding points in the latent space denoise the data, with trajectories that are smooth relative to the input data (Fig. 1E). Similar neural network approaches have been proposed to denoise from single-cell data, e.g. using a deep count autoencoder (Eraslan et al. 2019). RVAgene provides data denoising as a by-product of its primary functionality: modeling dynamic changes in gene expression.

Similar to other VAE-based tools for the analysis of single-cell data, RVAgene is efficient and scalable for use with large datasets. We compared the performance of RVAgene with a Bayesian nonparametric approach for the analysis of gene expression time series data (Dirichlet Process Gaussian Process (McDowell et al. 2018). Using either CPU or GPU computing, the time and memory gains are substantial, enabling the analysis of larger datasets than would otherwise be possible (Fig. S1). Analysis of RVAgene using simulated temporal data highlights the ability of such an architecture as means to study and generate gene expression dynamics. It enables learning of an unsupervised representation space, on which post-processing (e.g. unsupervised clustering) can be performed, as well as data denoising, and the generation of new time series data from arbitrary points in the latent space.

It is inevitably challenging to include sufficient dimensionality and variation in synthetic datasets to accurately capture biological processes such as those we observe in experimental datasets. Thus, in the subsequent two sections, we test the capabilities of RVAgene on two whole-genome biological datasets: embryonic stem cell differentiation, and kidney injury response. As we will see, in these cases it may not be possible to characterize the latent space by simple (e.g. k-means) clustering; we need to use other means to gain insight into the features of the latent space.

### 3.2 Characterization of embryonic stem cell differentiation with RVAgene

We applied RVAgene to model gene expression dynamics during embryonic stem cell (ESC) differentiation. Klein et al. (2015) identified 732 differentially expressed genes over the time course of mouse ESC differentiation following leukemia inhibitory factor (LIF) withdrawal. Data is gathered at four time points: 0, 2, 4, and 7 days after LIF withdrawal. (Table S2 in Klein et al. (2015)). We ordered the data (2717 single cells) using diffusion pseudotime (DPT), which provides robust methods for the reconstruction of single-cell temporal processes (Haghverdi et al. 2016). The root cell was randomly sampled from the initial time point (Fig. 2A-C). The inferred pseudotime is highly correlated with the experimental time points, giving confidence that true biological processes are represented over the DPT pseudotime. The gene expression dynamics over pseudotime show considerable variability among cells. To smooth the data, we apply a moving window average, over windows of length 40, to give 68 time points after smoothing (Fig. 2A-C). We fit linear regression models to the smoothed pseudotime profiles of each gene (Fig. S2), and see that for the majority of genes the correlation coefficients are > 0.5 (Fig. 2D), with a clear distinction between the up- and down-regulated genes over pseudotime.

**Figure 2.**
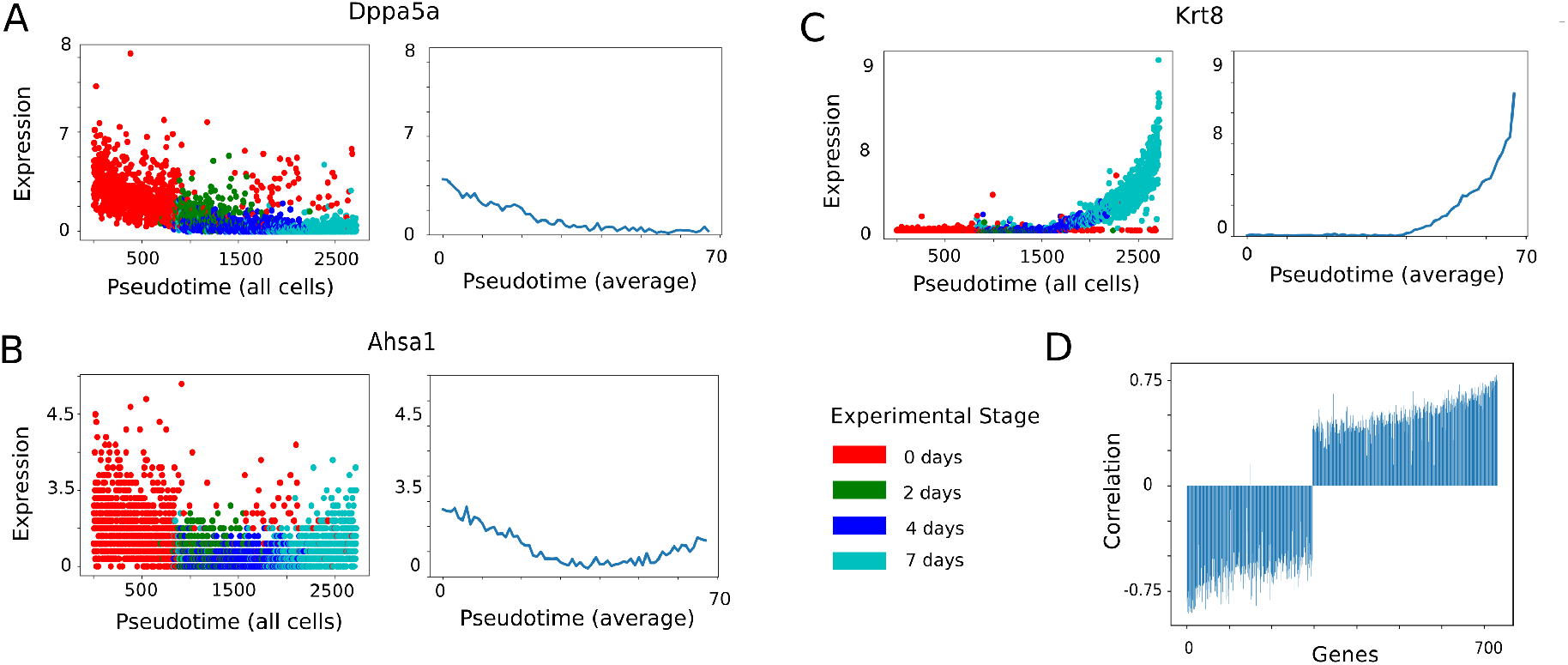
Analysis of embryonic stem cell differentiation gene expression dynamics in single cells. (**A-C**) Pseudotemporal ordering of 2717 single cells (data from (Klein et al. 2015)), calculated using DPT; three example genes shown: Dppa5a, Ahsa1, and Krt8. Gene expression values given as log2(counts+1) for all cells (left), and for sliding window average (right). (**D**) Pearson correlation coefficient between gene expression and time for 732 differentially expressed genes.

An RVAgene model was trained on the data with a two-dimensional latent space, on which genes are classified based on their correlation coefficients (Fig. 3A). Two distinctive characteristics emerge: a) the two groups (up- and down-regulated genes) are well-separated in the latent space, and b) the two groups merge and overlap at some point, illustrating the continuity of the latent space, as discussed above. We compared the results of RVAgene with DPGP, an unsupervised approach for gene expression time series clustering (McDowell et al. 2018). DPGP is a hierarchical Bayesian model that estimates the number of clusters along with the cluster membership, although to do so it has considerably higher resource consumption (Fig. S1). To assess the correspondence between methods, genes clustered by DPGP (Fig. S3) were projected onto the RVAgene latent space (Fig. 3B). Of the 12 clusters detected by DPGP, the four largest can be characterized by their up- and down-regulation profiles over pseudotime. On the RVAgene latent space, we find that genes sampled from each of the DPGP clusters appear close together, and moreover, are represented on a spectrum from upregulation to downregulation (Fig. 3B). The goals of RVAgene and DPGP are to some degree complementary: DPGP characterizes gene expression profiles discretely with no need for prior information, while RVAgene characterizes profiles with a continuous representation, that can explain smooth changes in patterns.

**Figure 3.**
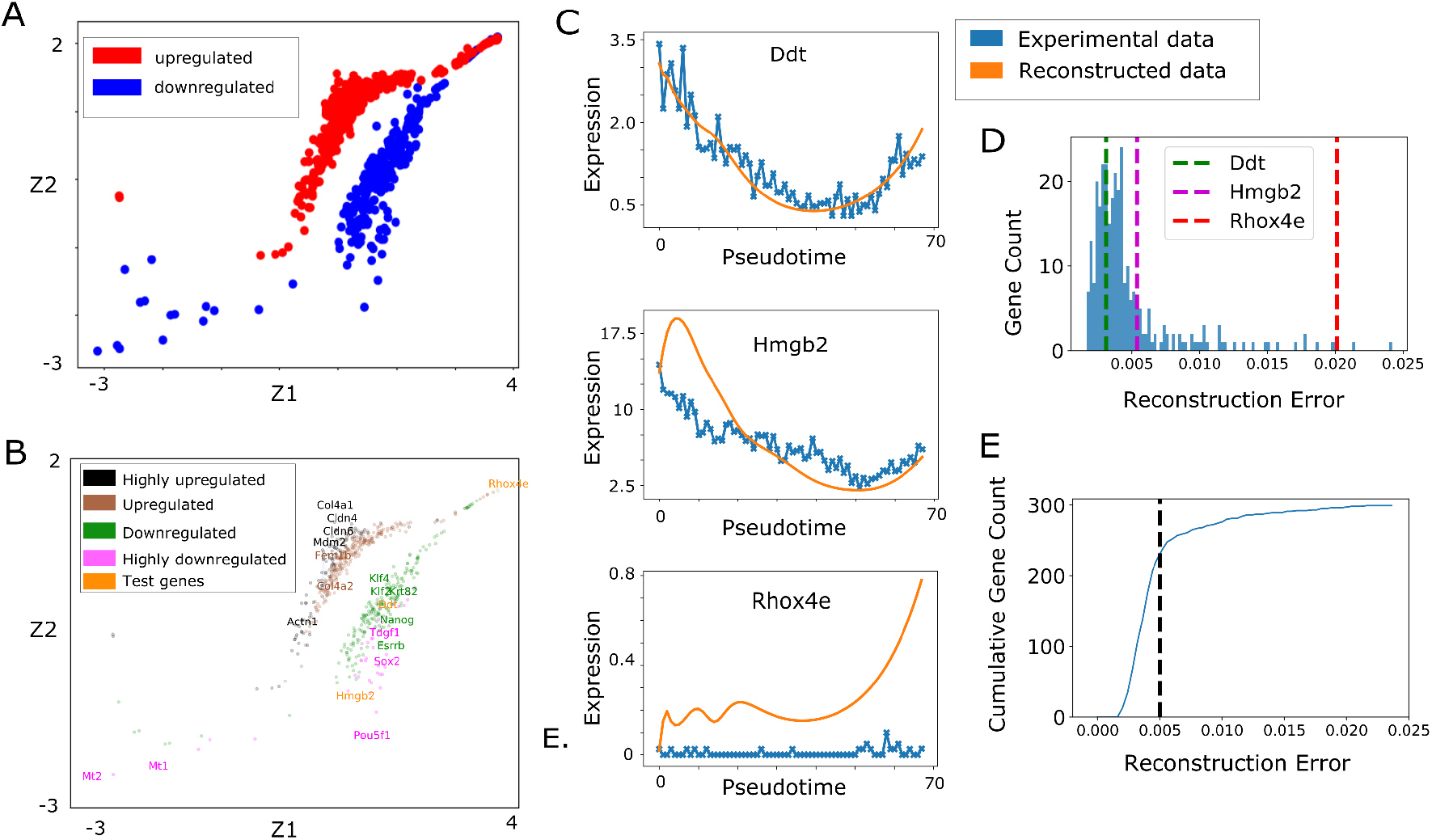
Accurate reconstruction of embryonic stem cell differentiation dynamics with RVAgene. (**A**) The 2D latent space learnt by an RVAgene model trained on 732 gene profiles over pseudotime, showing clear separation between upregulated and downregulated genes. (**B**) Comparison of RVAgene and DPGP. The four largest clusters from DPGP are plotted on the RVAgene latent space: temporal expression patterns (from highly upregulated to highly downregulated) are in close agreement between methods. (**C**) Comparison of experimental data and predictions. Model-generated reconstructions of three genes from the test set not used in training: Ddt, Hmgb2, and Rhox4e. Expression values are log2(counts+1). (**D**) Distribution of average *L*1 reconstruction errors for the 300 genes used in the test set. Genes plotted in C are marked. (**E**) Cumulative distribution of reconstruction error: 77% of genes (230/300) have reconstruction error less than 0.005.

To assess model predictions for individual genes, we kept aside 300 genes for testing and trained RVAgene on the remaining 432 genes. Predicted gene expression profiles are shown for three reconstructed genes, chosen to sample across the spectrum of reconstruction errors (Fig. 3C). The prediction for *Ddt*, which has a reconstruction error near the mode (Fig. 3D), shows very high accuracy. The prediction for *Hmgb2*, which has twice the reconstruction error, still broadly captures the temporal profile but with lesser accuracy. Finally we show the prediction for *Rhox4e*, a gene that was sampled from the long tail of the reconstruction error distribution, i.e. does not well match the data. Comparing these three examples with the full distribution of reconstruction errors (Fig. 3D), we see that the large majority of genes lie to the left of *Hmgb2*, i.e. have better-than-moderate prediction accuracy. The reconstruction error of *Hmgb2* is close to 0.005, which we use as a cut off for “well-reconstructed” genes, based on analysis of individual gene predictions. The cumulative reconstruction error distribution reiterates this point: 230 out of 300 genes (77%) have a reconstruction error ≤ 0.005 (Fig. 3E); we can conclude that the majority of test genes were faithfully reconstructed by the model.

RVAgene accurately reconstructed most gene profiles using only ~60% of the data for training (Fig. 4), likely due to co-regulation of gene expression programs. This led to a question: what is the smallest training gene set that can be used to accurately reconstruct gene dynamics? We subset the data randomly into train/test sets and trained separate RVAgene models on each. We found that reconstruction errors slowly increase as the size of the training set decreases, but not until the training set was as low as 18% of the data did the reconstruction errors significantly increase (Fig. 4). Analysis of the cumulative distribution of reconstruction errors across all groups found that RVAgene predicts the majority of gene temporal profiles well (defined as below a reconstruction error of 0.005) if ≥ 45% of the data is used for training. The successful prediction of gene expression dynamics de novo using small subsets of the data suggests widespread co-regulation of gene expression programs during embryonic stem cell differentiation, as found in previous work (Jang et al. 2017).

**Figure 4.**
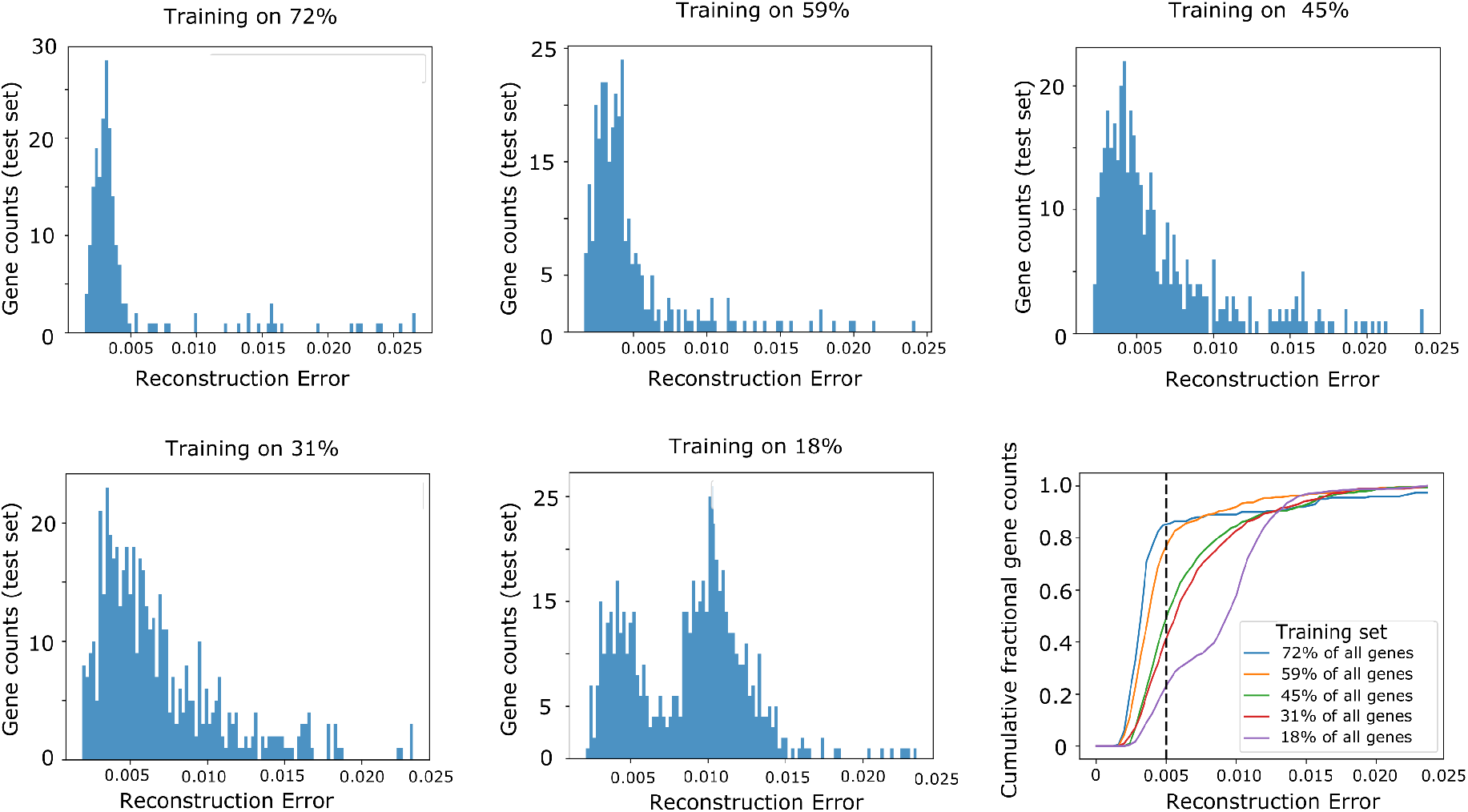
Accuracy of RVAgene reconstructions for different train/test group sizes. Distributions of reconstruction errors on randomly sampled sets of test genes, where the full data were split into test groups of: 200 genes (train on 72%), 300 genes (train on 59%), 400 genes (train on 45%), 500 genes (train on 31%), and 600 genes (train on 18%). Cumulative fractional distribution of reconstruction errors (cumulative count/test set size) for all groups.

### 3.3 RVAgene can classify and predict gene expression dynamics in response to kidney injury

We investigated gene expression dynamics in the murine kidney by applying RVAgene to a dataset that describes gene expression profiles before, during, and after a kidney injury (Liu et al. 2017). The dataset is temporally rich, with a total of ten bulk samples over twelve months. Since in this case no single-cell information is available, we cannot order samples by pseudotime to smooth the data. Moreover, the temporal gene expression profiles described in Liu et al. (2017) display more complex dynamics than for the previous dataset (Klein et al. 2015), and are not readily separable by linear patterns of up- and down-regulated genes (cf. Fig. 3A). Thus, below, we must consider nonlinear models in order to characterize the temporal patterns observed.

The data consist of one initial timepoint (*t* = 0) before the injury event (an ischemia/reperfusion injury model) and nine subsequent time points (*t* = 1 to 10) following the injury (48 hours, 72 hours, 7 days, 14 days, 28 days, 6 months and 12 months). We note that the timepoints are not uniformly spaced, which is not taken into account in RVAgene, which only models the broad temporal trend (see Discussion). From an initial list of 1927 differentially expressed genes measured over the time course in three biological replicates, we removed putative/predicted and non-protein coding genes, retaining a list of 1713 genes as input to the model.

We ran RVAgene separately for each of three biological replicates. Independent replicates & independently trained models provide additional means with which to test the reproducibility of these methods. For each replicate, RVAgene was trained with a two-dimensional latent space and a hidden size of 10, on the full set of genes over 200 epochs: found to be sufficient for the convergence of 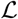 (see Methods for further details). We fit linear regression models to the temporal gene profiles (Fig. S4) and found that linear fits rarely described the gene temporal profiles well (most correlation coefficients had values close to zero), not did they identify separate clusters in the latent space. Normalizing the data to lie in [0, 1] improved our ability to discriminate clusters in the latent space (Fig. S4C), but came at the expense of a significant loss of information, as the variance captured in the latent space was dramatically reduced. The absence of evidence for linear correlations could indicate expression dynamics that are uncorrelated with time, but could of course also indicate more complicated (nonlinear) gene expression dynamics, which are explored below.

To study nonlinear gene expression dynamics, we fit a 2nd degree polynomial, i.e. we fit the temporal trajectory of each gene *x* to: *x* = *at*^2^ + *bt* + *c*, where *a, b, c* are constants (Fig. S5). We hypothesized that this function could adequately describe the transient dynamics observed by Liu et al. (2017) for most genes in response to the kidney injury. Thus, we classified genes into one of two groups, *a* < 0: convex (up-down pattern), 1200 genes; and *a* ≥ 0: concave (down-up pattern), 512 genes. In the latent space, the separation of these two groups is clearly visible for each replicate (Fig. 5A). Moreover, the classification is in agreement with Liu et al. (2017), where the majority of differentially expressed genes are upregulated transiently. To explore the ability of RVAgene to generate gene expression dynamic profiles de novo, we kept aside 300 randomly sampled genes for testing, and trained RVAgene models on the remaining genes for each of the three replicates. Independently for each model, we then generated dynamic profiles for the test genes. Three genes sampled randomly from the test set are plotted in Fig. 5B. Of particular note, for each of genes, the model-generated data captures the temporal patterns while displaying a higher degree of similarity across replicates than the experimental data itself. This illustrates that the model is neither under-nor overfitting, but capturing the underlying biological patterns while sufficiently accounting for the noise. Reconstruction errors are comparable across the three replicates, albeit with slightly higher overall errors in replicate 1 (Fig. 5C-D). Overall, the reconstruction errors are higher than for the previous section (averaging over many pseudotemporal time points allowed us to significantly reduced the noise).

**Figure 5.**
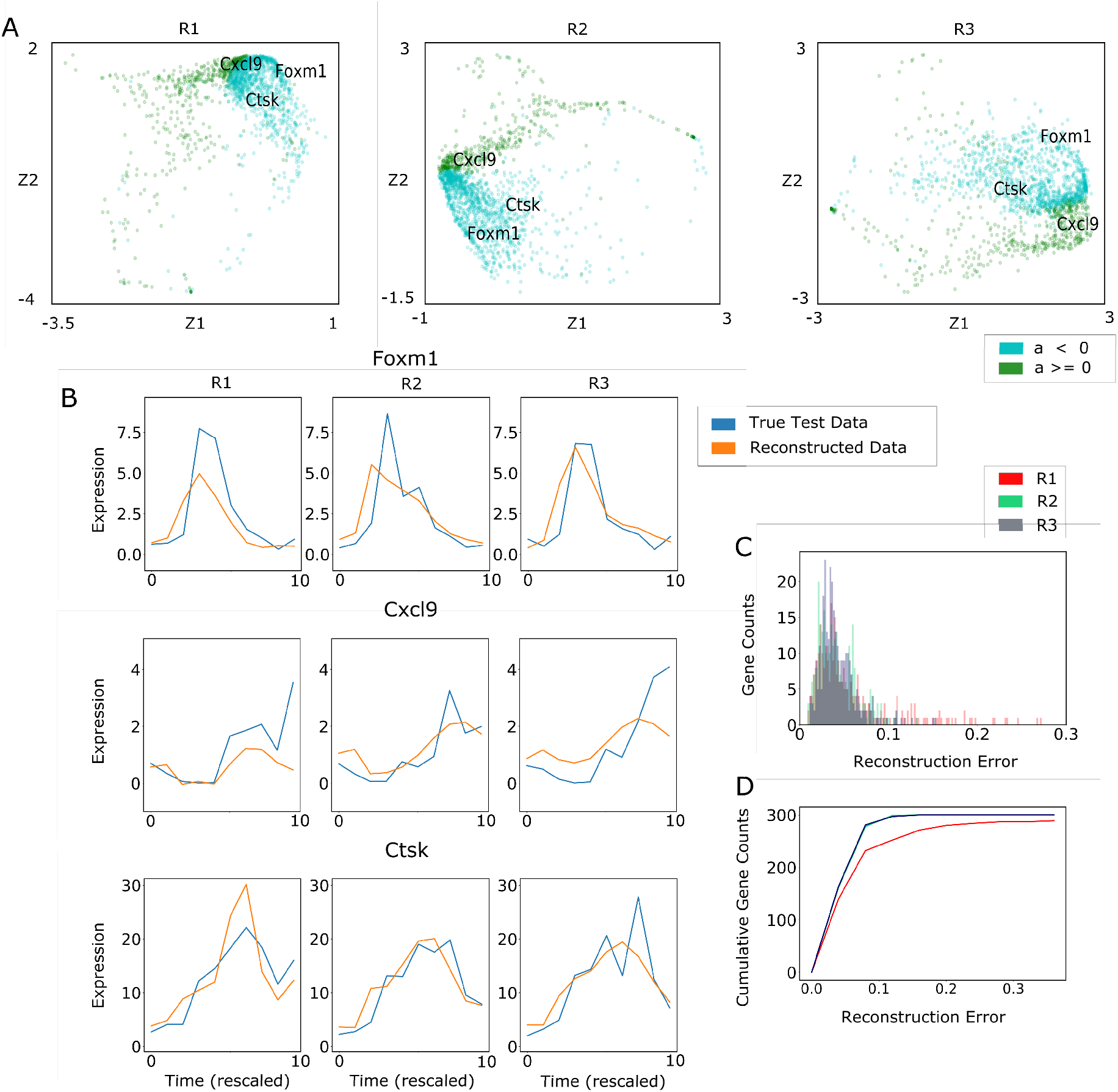
Accurate reconstruction of kidney injury response gene dynamics with RVAgene. (**A**) Latent space representations of RVAgene models trained separately on three independent replicates (R1-R3); classified by quadratic fit coefficient *a*. (**B**) Model generation of gene dynamics for genes not used in training: *Foxm1, Cxcl9* and *Ctsk*. (**C**) Histograms of reconstruction errors for RVAgene models trained on R1-R3 (truncated). (**D**) Cumulative distribution of reconstruction errors.

To investigate in more depth the features that are captured in the RVAgene latent space, we studied three distinct areas of the latent space of each model. Areas were defined using three gene groups, chosen simply based on their co-location in distinct regions of the latent space: 1) a *Wnt* group consisting of family members *Wnt2* & *Wnt4*; 2) an *Slc* group consisting of family members *Slc7a13* & *Slc22a18*; and 3) a *Sdc1* group, consisting of only *Sdc1*. For each group, we characterized neighboring genes by defining a circular neighborhood around each gene in the group, with radius *r* (depending on the local density, the radius was varied, giving: *r*^2^ = 1 for *Slc*, *r*^2^ = 0.3 for *Sdc*, *r*^2^ = 0.05 for *Wnt*. We then took all genes inside this radius for each replicate, and found the intersection of genes over the three replicates (Fig. 6A-B). We analyzed the intersection gene set for each group by studying their temporal profiles and their gene ontology (GO) term associations. Each group was characterized by a strikingly clear temporal profile. The *Sdc1* and *Wnt* groups both show transient upregulation, over different timescales: the *Sdc1* group is upregulated from 24 hours post-injury until 14-28 days post-injury (fast response) (Supplementary Fig. S6B), whereas the *Wnt* group is upregulated at 7 days post-injury until 28 days post-injury (slow response) (Fig. 6C). In contrast, the *Slc* group is downregulated at 24 hours post-injury, and remains suppressed until 7-28 days post-injury (Fig. 6D).

**Figure 6.**
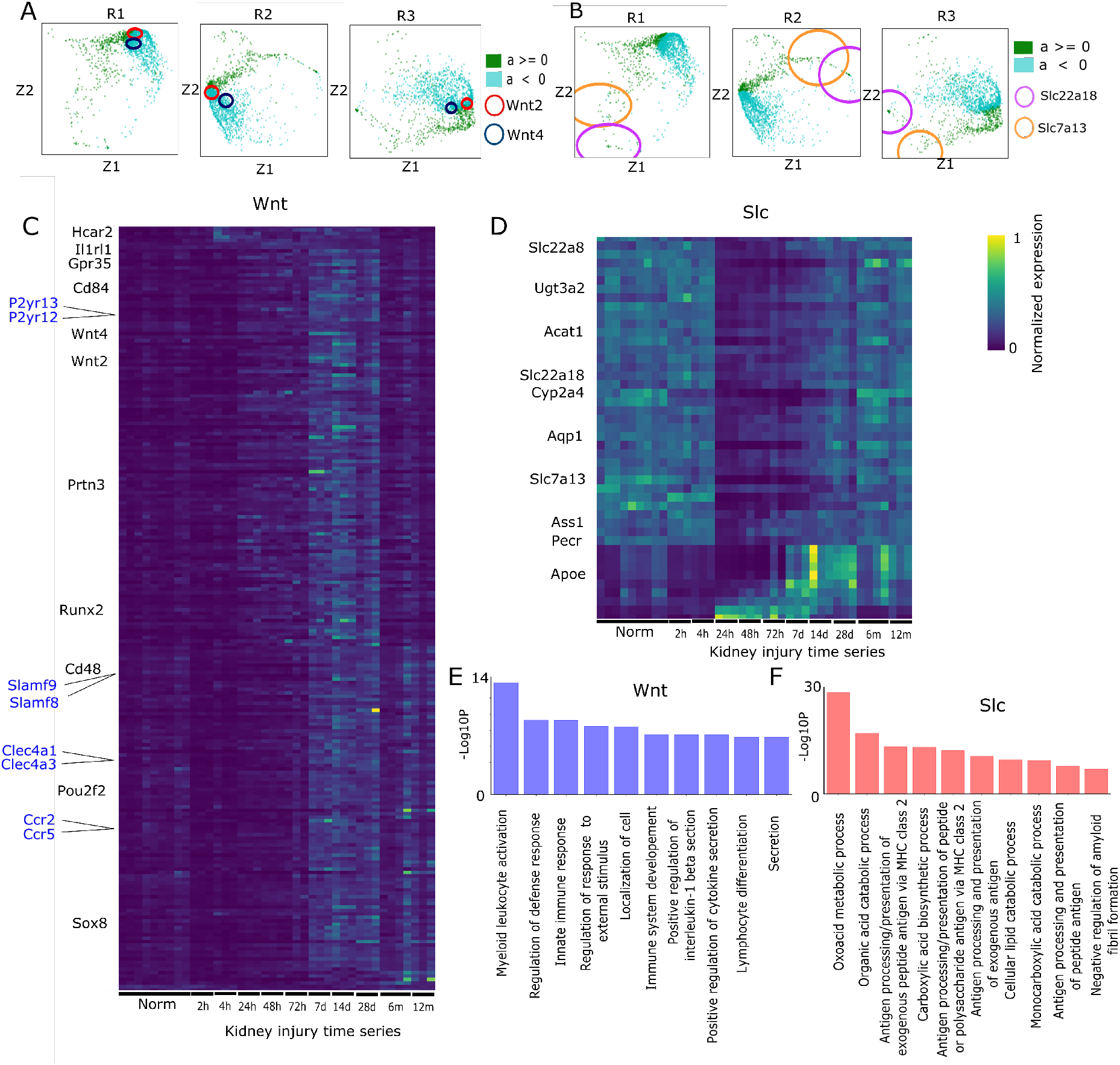
RVAgene latent space captures biological processes driving concordant gene expression hanges. (**A**) Z-plots for replicates R1-R3 with local neighborhoods of Wnt2 and Wnt4 marked (circles). (**B**) As in A, for Slc family members Slc22a18 and Slc7a13. (**C**) Heatmap of expression changes over time course of injury for the Wnt neighborhood genes in the intersection of R1-R3. Selected genes marked (black), as well as ortholog gene pairs (blue). (**D**) As in C, for Slc neighborhood genes. (**E**) Histogram of −log10 p values of gene ontology terms for biological processes terms associated with the Wnt neighborhood (gene set in C). (**F**) As in E, with the Slc neighborhood (gene set in D).

Analysis of GO biological process terms enriched in each gene group further highlighted the power of the latent space for biological discovery. The fast response (*Sdc1*) group was characterized by upregulation of programs related to apoptosis, stress response, wound healing and chemotaxis, i.e. the first responders to the site of injury (Fig. S6C). In addition all five *Lox* genes comprising the GO term “peptidyl-lysine oxidization” were found in this group. This is consistent with the oxidative stress resulting from the renal ischemia-reperfusion injury that was performed. However, distinct factors regulate the *Lox* family genes, as can be partly observed by their subtle differences in temporal profile (Fig. S6D). Their co-location in the latent spaces of all three models thus highlights the potential use of RVAgene for discovery of complex temporal regulatory events from gene expression data.

The slow response (*Wnt*) group was primarily characterized by immune response processes, including leukocyte activation, platelet aggregation, and various cytokine-mediated pathways including *IL-1* and *IL-33* (Fig. 6E). Notably, the Wnt group identies multiple gene orthologs (Fig. 6C) with very similar profiles: likely evidence of shared temporal regulation. This illustrates once again (as for the *Lox* genes above) the potency of RVAgene for the discovery of temporally co-regulated genes.

Finally, the *Slc* group of genes shows a transiently down-regulated pattern between 24 hours and 7-28 days, although some gene in this group deviate from this pattern (Fig. 6D). GO term enrichment identifies the positive regulation of metabolic processes (Fig. 6F). The downregulation of metabolic programs during the response to kidney injury is agreement with the findings of Liu et al. (2017). Notably, this metabolism-sensitive group contains many genes that also display sexually dimorphic expression, primarily in specific regions of the proximal tubule (Ransick et al. 2019), thus independently identifying the well-established (though under-studied) interplay between sex differences and injury responses in the kidney (Neugarten et al. 2000).

In summary, unsupervised analysis of groups of genes co-located in the latent spaces of RVAgene finds: 1) high similarity between temporal gene profiles of genes nearby in latent space, and 2) clear biological signatures represented by these groups of nearby genes, in strong agreement with prior knowledge (Liu et al. 2017). Moreover, the latent spaces of RVAgene models can be used to predict programs of temporal co-regulation.

## 4 Discussion

We have presented RVAgene, a recurrent variational autoencoder for generative modeling of gene expression time series data. Through its encoder network, RVAgene provides means to visualize and classify gene expression dynamic profiles, which can lead to the discovery of biological processes. Through its decoder network, RVAgene provides means to generate new gene expression dynamic profiles by sampling points from the latent space, and can accurately predict gene dynamics in complex biological data. As a by-product, the model produces smoothed outputs, which can be used for denoising gene expression time series data. RVAgene is efficient on temporally-rich whole genome datasets, in comparison to current methods used (Gaussian processes).

RVAgene can be used to discover structure in the data, such as gene profile clusters. Popular methods for clustering gene profiles such as Bayesian hierarchical clustering (Cooke et al. 2011) or DPGP (McDowell et al. 2018) detect the number of clusters in the data by fitting a hyperparameter *α*, the concentration parameter of the governing Dirichlet process (Ferguson 1973). Although unsupervised, inevitably, the choice of *α* affects the number of clusters output. Visualizing the data first with RVAgene can give an idea whether the data favor clustering or a continuous representation. Thus analysis in RVAgene can guide the setting of the hyperparameter *α* in DPGP and similar methods. In the case of ESC differentiation, DPGP predicts 12 clusters (Fig. S3), yet most have very few members and many share similar patterns. The RVAgene latent space for this dataset finds two major divisions in the data, and orders the largest DPGP clusters along a spectrum (Fig. 3B), suggesting that DPGP might be overfitting the data. Indeed, the two methods can be used complementarily: RVAgene for high-level structure discovery and DPGP for clustering. We note however that DPGP does not scale well with large datasets and thus cannot always be used (Fig. S1).

The latent space of an RVAgene model encodes useful information about biological features, and in that sense provides biologically interpretable representations of the data. However, the representation is not interpretable in the sense that the components of the latent space do not have a physical meaning nor are they necessarily independent. Recent methods have tackled this issue of interpretability, by either modifying the loss function to make components independent (Higgins et al. 2016) or substituting linear functions in parts of the VAE (Svensson et al. 2020, Ainsworth et al. 2018). These methods have clear advantages regarding the analysis and interpretation of features in the latent space. In future work, decoding an RVAgene model with a linear function (Svensson et al. 2020) could facilitate additional discovery and improve our ability to gain insight into dynamic biological processes through the analysis of the latent space.

Dynamic changes in gene expression underlie essential cell processes. As such, modeling gene expression changes can also facilitate downstream analysis tasks, including gene regulatory network (GRN) inference. Inferring gene regulatory networks from single-cell data is challenging (Chen & Mar 2018), particularly due to cell-cell heterogeneity and high levels of noise. Several recent approaches to GRN inference use differential equations in their formulation to model gene expression changes (Ma et al. 2020, Aubin-Frankowski & Vert 2020, Matsumoto et al. 2017). RVAgene could either supplement such methods directly by providing denoised input data, or could be used to replace the differential equation-based components of these methods (which are notoriously difficult to parameterize) with a generative model of gene expression dynamics.

RVAgene is currently agnostic of irregular time intervals between consecutive points in a time series, i.e. it standardizes the time interval. This is not usually a concern for single-cell data, since with pseudotime information we can choose appropriate time intervals. However, in other cases, such as in response to kidney injury (Liu et al. 2017), standardizing time intervals distorts the dynamic profiles. Since RVAgene seeks to describe broad temporal patterns, we do not see this as a critical issue, though it would be desirable to generalize the model. A simple way to model irregularly spaced time points would be to augment the data through interpolation, though this is difficult without making strong assumptions about the (generally unknown) noise model. Gaussian process models (McDowell et al. 2018, Hensman et al. 2013) can take irregular data as input, although (as noted above) are not efficient enough to run on large datasets. An alternative approach would be to modify the recurrent network architecture to take time points explicitly as input values, this would enable modeling of irregular or asynchronous data (Wu et al. 2018).

RVAgene models in discrete time steps. There is no simple modification to the recurrent network structure that allows for prediction on continuously valued time. However, a recent development: neural ordinary differential equations (ODEs) (Chen et al. 2018), enables modeling of time series data with continuous timepoints. Chen et al. (2018) describe a generative latent ODE architecture similar to that of RVAgene, except that in their case the recurrent decoder network is replaced by a neural ODE decoder network. Chen et al. (2018) demonstrate accurate results using synthetic data, however when we applied the method to the ESC single-cell differentiation dataset (Klein et al. 2015), the neural ODE network was found to converge very slowly and was overall underfit (Fig. S7). The latent ODE method used by Chen et al. (2018) does not address the challenge of modeling asynchronous/irregularly spaced data, but this has been more recently addressed (Rubanova et al. 2019). These new models may well lead to future improvements in network architectures, although it seems that computational progress is needed before they can be successfully applied to complex biological systems.

In the current work, the prior on latent space used throughout was a unit spherical Normal, appropriate for exploratory data analysis where we have no further knowledge about structure in the latent space. However, given more information, e.g. that the data is contains *k* clusters, a different prior on the latent space might be more appropriate. A multi-modal prior – such as a Gaussian Mixture Model (GMM) prior – would permit structured (multi-modal) representations. However, the KL-divergence for an arbitrary GMM is not tractable; approximation (Hershey & Olsen 2007) or numerical computation would be necessary. Moreover, there is a greater problem: mixture models contain discrete parameters and VAE models are ill-suited for the optimization of discrete parameters (Dilokthanakul et al. 2016), thus directly replacing the Normal prior of a VAE with a GMM is not feasible. A workaround to this problem is presented in (Dilokthanakul et al. 2016), however implementing this for a recurrent model architecture remains an open problem.

The points raised above offer much scope for future work. These include the design of new latent space models with informative priors, modeling irregular time series data, and modeling in continuous time. Developments in some of these areas (Chen et al. 2018), while promising, tend to rely on training data with relatively low levels of noise: far from the reality of most biological data. Thus it seems highly likely to be beneficial for both machine learning and biology to develop new neural network architectures in light of biological data.

## Aknowledgements

We thank A.P. McMahon for valuable discussions and comments on the manuscript. R.M. gratefully acknowledges support from a USC Viterbi Fellowship.

## Data Availability

The synthetic data used for evaluation of RVAgene are available at: https://github.com/maclean-lab/ RVAgene. Additional data used in the manuscript are available from the Gene Expression Omnibus: ESC differentiation (GEO accession GSE65525) and kidney injury (GEO accession GSE98622).

## Software Availability

RVAgene is available in Python released under an MIT license: https://github.com/maclean-lab/RVAgene.

## Supplementary Information

Supplementary figures are available for download.

