## Supplemental Figures for "RVAgene: Generative modeling of gene expression time series data"

### Recurrent Variational Autoencoder for Generative Modelling of Gene Expression Time-Series Data

Raktim Mitra, Adam L. MacLean*
**Quantitative and Computational Biology, University of Southern California**
*****

#### Supplementary figures

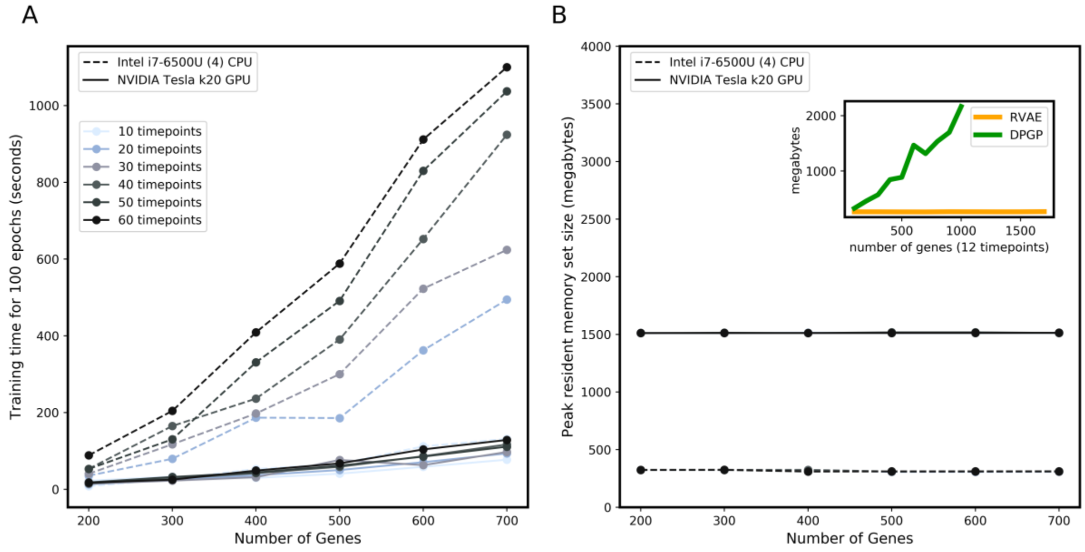


**Figure S1. Training RVAgene is scalable on CPU and GPU.** (**A** ) Computational cost of training RVAgene for 100 epochs of training with varying number of genes and time points on an intel i7 CPU and Tesla K20 GPU. As the number of time points and genes grows large (up to 60 time points and 700 genes tested), total runtimes on CPU were on the order of $10^3$ seconds ($< 20$ minutes). Runtimes on a GPU were reduced around 100 seconds ($< 3$ minutes). Thus RVAgene is scalable to tens of thousands of genes and hundreds of time points for training times up to some days on a CPU but only hours on a GPU. (**B**) In terms of peak memory usage, for RVAgene (a neural network), this is defined by the backpropagation, and is constant (proportional to the size the network). In comparison, for DPGP [McDowell et al., 2018], memory storage requires $G$ matrices of size $T \times T$, where $G$ is number of genes in the data and $T$ is number of timepoints per gene: for large datasets, this quickly becomes infeasible (inset plot shows runtime peak resident set size). On the ESC differentiation data (732 genes and 68 time points) [Klein et al., 2015], it was not possible to run DPGP on a machine with 8 CPU cores 16 GB RAM. Only by reducing the number of time points to 13 (by time-averaging) could DPGP be run without memory error.

1

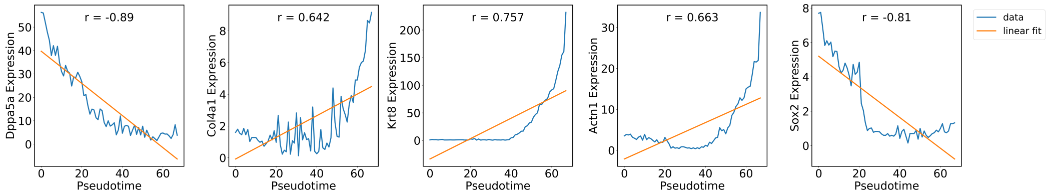


**Figure S2**. Characterization of gene dynamics by linear fit using Pearson correlation coefficient for 5 sample genes in the ESC differentiation dataset [Klein et al., 2015]. Blue lines represents original data and orange lines represents linear fits. The Pearson correlation coefficient $r$ is given for each plot.

2

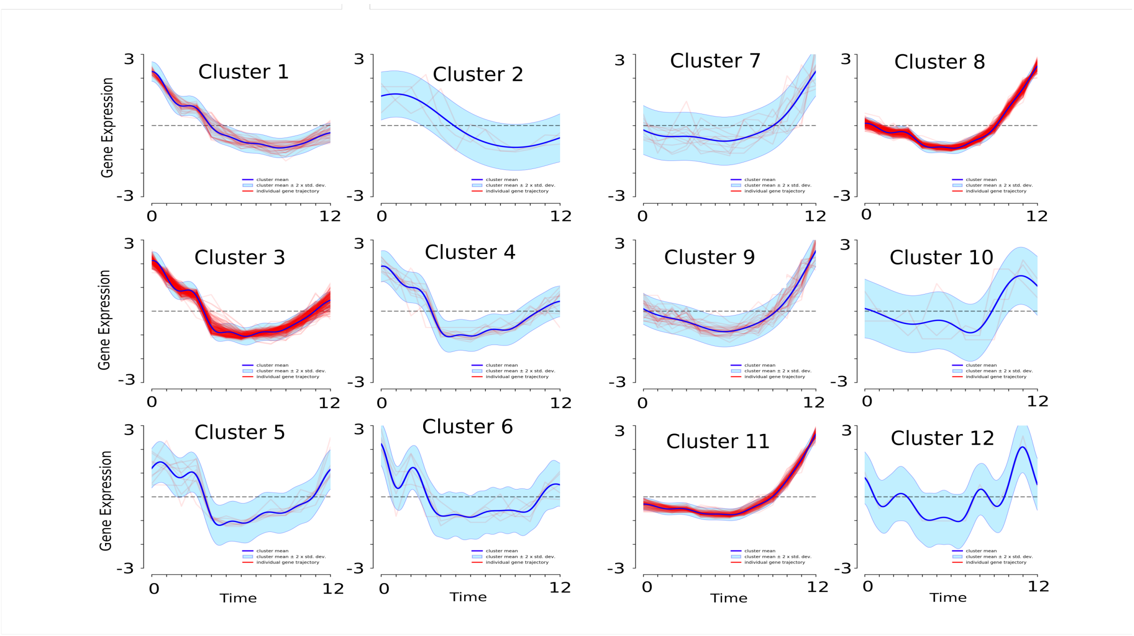


**Figure S3. Clusters detected by the unsupervised clustering algorithm DPGP for ESC differentiation.** Clusters detected by DPGP in the ESC differentiation dataset [Klein et al., 2015] with default hyperparameters showing cluster means (black), mean $\pm$ 2 s.d. in (blue) and cluster members (red).

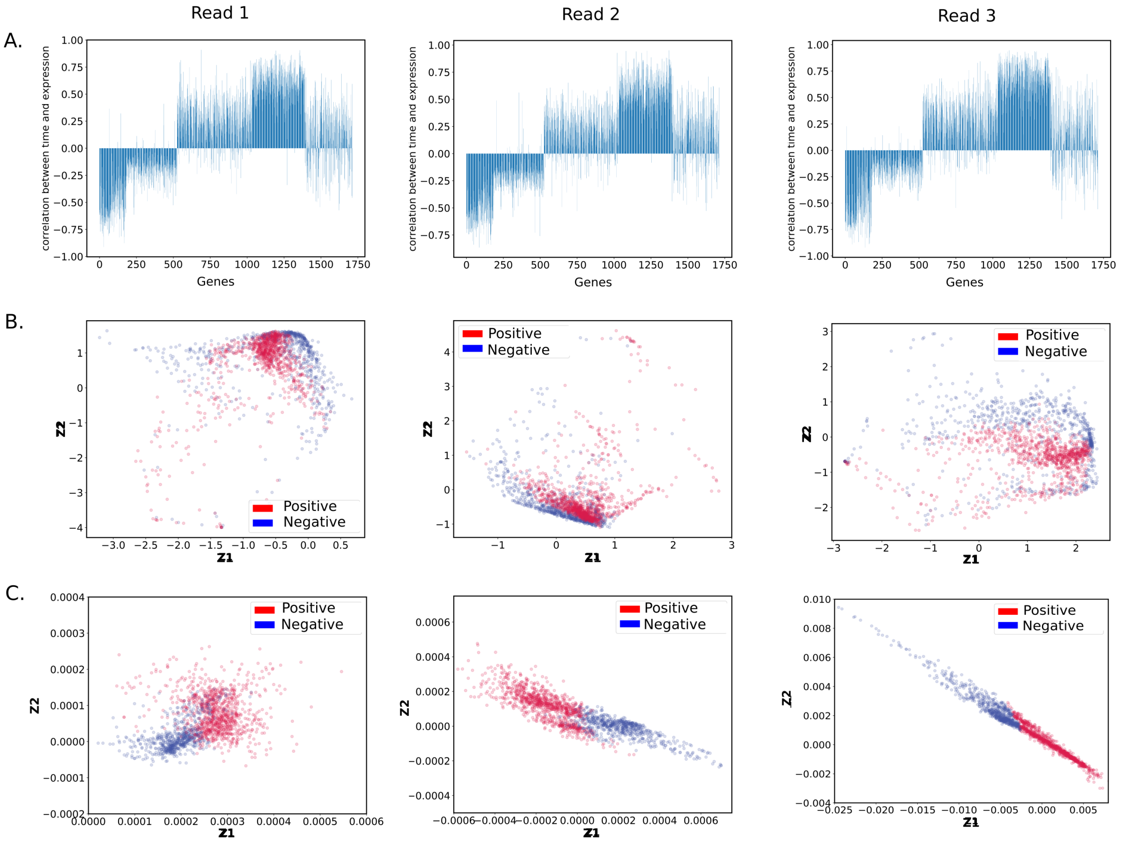


**Figure S4. Modeling response to kidney injury and analysis of linear fits.** (**A**)) Pearson correlation coefficients between gene expression and time for each differentially expressed gene in the kidney injury dataset for each of the 3 replicates [Liu et al., 2017]. (**B**)) RVAgene latent space representation of fitted model for each replicate; color represents positive or negative correlation coefficients. (**C**)) RVAgene latent space representation learnt for the same three replicates as in (B), but where every input gene was normalized so that its expression sums to 1.

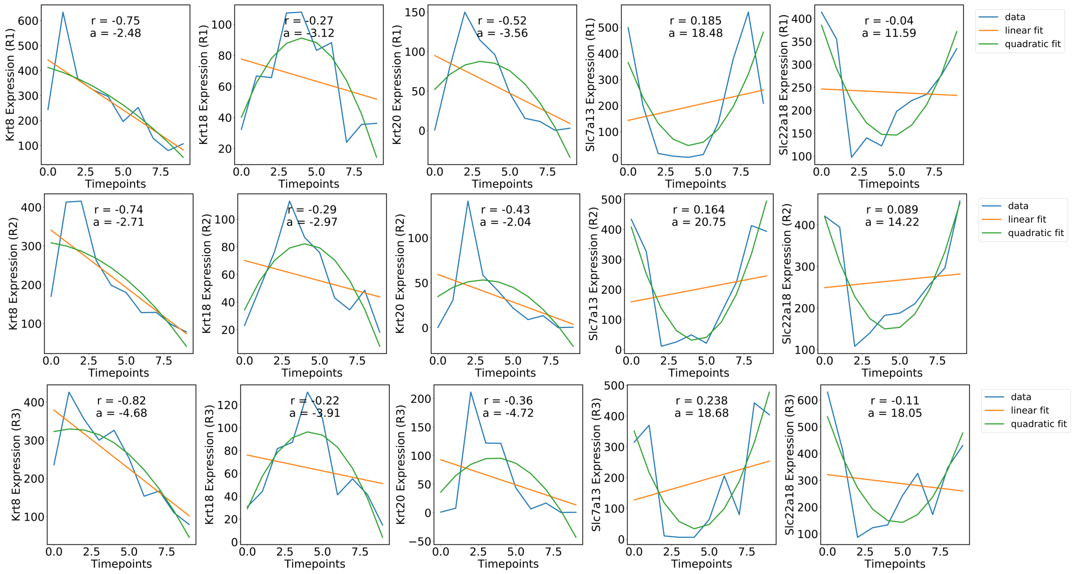


**Figure S5. Comparison of linear and quadratic fits to describe gene dynamics in response to kidney injury.** For each of the three replicates (R1-R3), five genes are shown, with experimental data (blue), linear fit (orange), and quadratic fit (green). Pearson correlation coefficients, $r$, and quadratic coefficients, $a$ ($x = at^2 + bt + c$), are given for each plot.



5

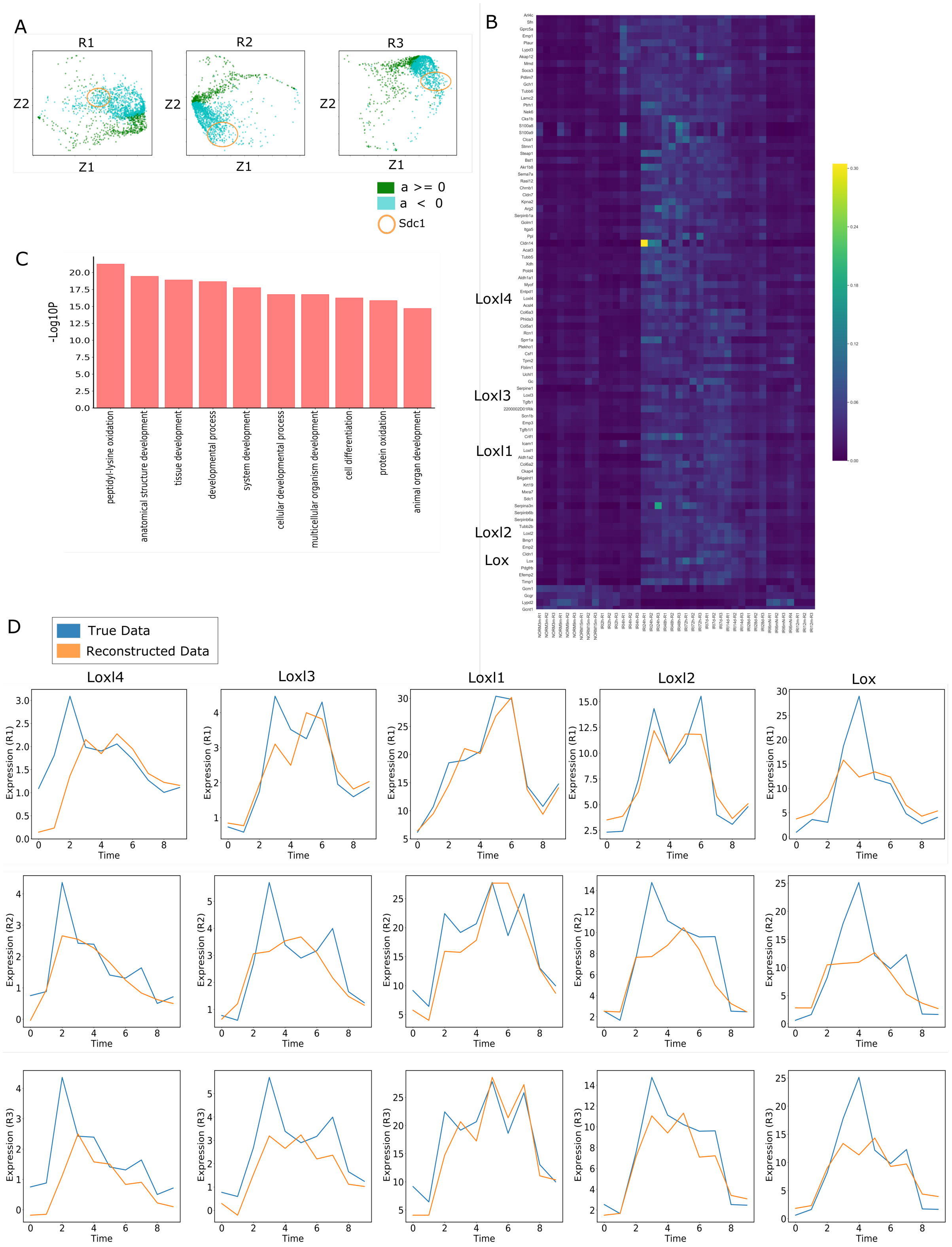


**Figure S6. RVAgene latent space captures biological processes driving concordant gene expression changes. (A)** Latent space representations for replicates R1-R3 with local neighborhoods of Sdc1 marked (circles). **(B)** Heatmap of expression changes over time course of injury for the Sdc1 neighborhood genes in the intersection of R1-R3; selected genes highlighted. **(C)** Histogram of -log10 p values of top gene ontology terms for biological processes terms associated Sdc1 neighborhood genes (gene set in B). **D)** Predicted vs true data plotted for each of the Lox genes identified in (B), for each of the three replicates.

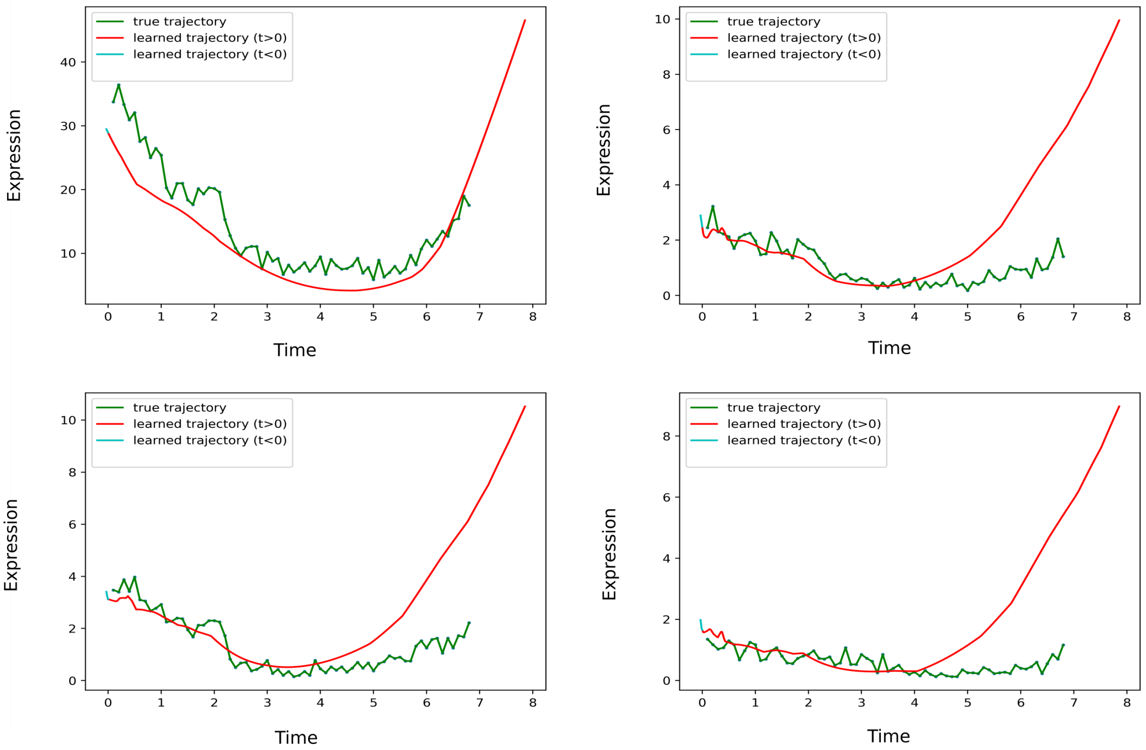


**Figure S7. Examples of continuous time prediction of ESC differentiation**. Reconstruction (up to $t = 6.8$) and future prediction (for $t > 6.8$) for 4 example genes by a latent ODE [Chen et al., 2018] trained on ESC data [Klein et al., 2015] for 1000000 iterations, showing a good fit for the initial timepoints, but underfitting for the later timepoints.

7
